# XBB.1.5 monovalent booster improves antibody binding and neutralization against emerging SARS-CoV-2 Omicron variants

**DOI:** 10.1101/2024.02.03.578771

**Authors:** Shilpi Jain, Sanjeev Kumar, Lilin Lai, Susanne Linderman, Ansa A. Malik, Madison L. Ellis, Sucheta Godbole, Daniel Solis, Malaya K. Sahoo, Kareem Bechnak, Isabel Paredes, Ralph Tanios, Bahaa Kazzi, Serena M. Dib, Matthew B Litvack, Sonia T. Wimalasena, Caroline Ciric, Christina Rostad, Richard West, I-Ting Teng, Danyi Wang, Sri Edupuganti, Peter D. Kwong, Nadine Rouphael, Benjamin A. Pinsky, Daniel C. Douek, Jens Wrammert, Alberto Moreno, Mehul S. Suthar

**Author notes:** Co-corresponding authors: Mehul S. Suthar; Alberto Moreno.

## Abstract

The rapid emergence of divergent SARS-CoV-2 variants has led to an update of the COVID-19 booster vaccine to a monovalent version containing the XBB.1.5 spike. To determine the neutralization breadth following booster immunization, we collected blood samples from 24 individuals pre- and post-XBB.1.5 mRNA booster vaccination (∼1 month). The XBB.1.5 booster improved both neutralizing activity against the ancestral SARS-CoV-2 strain (WA1) and the circulating Omicron variants, including EG.5.1, HK.3, HV.1, XBB.1.5 and JN.1. Relative to the pre-boost titers, the XBB.1.5 monovalent booster induced greater total IgG and IgG subclass binding, particular IgG4, to the XBB.1.5 spike as compared to the WA1 spike. We evaluated antigen-specific memory B cells (MBCs) using either spike or receptor binding domain (RBD) probes and found that the monovalent booster largely increases non-RBD cross-reactive MBCs. These data suggest that the XBB.1.5 monovalent booster induces cross-reactive antibodies that neutralize XBB.1.5 and related Omicron variants.

## Main

The development and fast deployment of the first generation of COVID-19 vaccines has changed the course of the pandemic by protecting against severe disease outcomes that resulted in hospitalization and death. The World Health Organization estimates that as of December 8, 2023, over 13.6 billion vaccine doses have been administered worldwide. The vaccination efforts during the first year of implementation are estimated to have averted 19.8 million deaths globally^1^. The continuous evolution of severe acute respiratory syndrome coronavirus 2 (SARS-CoV-2), the rapid emergence of antigenically divergent variants, and waning vaccine effectiveness against SARS-CoV-2 infection has led to updates in the COVID-19 vaccine formulations. Bivalent mRNA COVID-19 vaccines have been available since September 2022 and cover variants that are no longer circulating^2^. In September 2023, the United States Food and Drug Administration (FDA) has approved a monovalent XBB.1.5 vaccine to protect against circulating variants^3^. The monovalent mRNA vaccine includes only the XBB.1.5 spike and aims to improve protection against contemporary Omicron variants.

The introduction of booster immunizations was implemented to counteract waning immunity and promote humoral responses against emerging variants^4,5^. Both primary mRNA vaccine series and a third-dose immunization protected against severe disease, but vaccine effectiveness against infection declined significantly with the emergence of the Omicron variants^6,7^. We have shown that a booster third dose has a minor effect on increasing the half-life of antibody titers and showed no effect on inducing neutralizing antibodies against novel Omicron variants^5^. With the introduction of bivalent booster mRNA vaccines in September 2022 that include the ancestral WA1 and BA.5 spike, the neutralizing antibody response elicited exhibited a greater breadth^8^. However, Omicron variants that circulated early in 2023, such as BQ.1.1 and XBB.1, demonstrated an ability to escape neutralizing antibodies^9–11^, presumably contributing to reduced vaccine effectiveness^12^. Since the FDA approval of the updated monovalent XBB.1.5 vaccine in September of 2023, rapid evolution of additional subvariants has been identified. The JN.1 variant, which is related to the BA.2.86 variant, contains an additional spike mutation at position L455S and has rapidly become the dominant variant in the United States and throughout the world^13^. Further, HV.1, which is closely related to XBB.1.5 and EG.5.1, currently contributes to approximately 5% of total COVID-19 cases in the United States.

## Results

### XBB.1.5 monovalent booster improves neutralizing activity against emerging Omicron variants

We obtained plasma samples from 24 participants before and after XBB.1.5 monovalent vaccine administration (13-41 days post-booster dose). All individuals received three to four doses of ancestral mRNA monovalent booster followed by either none or one dose of bivalent vaccine (**Supplementary Table S1**). We used the FRNT in a VeroE6/TMPRSS2 cell line to compare the neutralizing activity against the WA.1 isolate of SARS-CoV-2, B.1.617.2 (Delta), and Omicron subvariants, BA.5, EG.5.1, XBB.1.5, HK.3, HV.1, and JN.1 in pre- and post-boost samples of these individuals. FRNT_50_ (the reciprocal dilution of serum that neutralizes 50% of the input virus) and geometric mean titers (GMTs) were calculated. Serum samples in which the GMT fell below the limit of detection (1:20) were given an arbitrary FRNT_50_ value of 10, and these samples are considered as undetectable or non-responders against the respective variant.

In all the individuals before the XBB.1.5 immunization, neutralizing activity was lower against all Omicron subvariants than against the WA1 strain, with a reduction in neutralization potency more than 6-fold for BA.5, 14-fold for EG.5.1, 29-fold for XBB.1.5, 25-fold for HK.3, 24-fold for HV.1, and 19-fold for JN.1. Neutralizing activity was lowest against the XBB.1.5 subvariant. The FRNT geometric mean titers (GMTs) were 650 for WA1, 390 for B.1.617.2, 116 for BA.5, 47 for EG.5.1, 22 for XBB.1.5, 26 for HK.3, 27 for HV.1, and 34 for JN.1. All participants had detectable neutralizing antibodies to the WA1 and B.1.617.2 viruses, while 88% of the individuals made nAb to BA.5 and 75% had nAb titers to EG.5.1 variant. Only 54% of individuals (13/24) have detectable neutralizing antibodies against XBB.1.5 variant before vaccination.

Neutralizing antibodies against the newly circulating Omicron variants HK.3, HV.1, and JN.1 were undetectable in around 30% of individuals (**Fig. 1a**).

**Figure 1.**
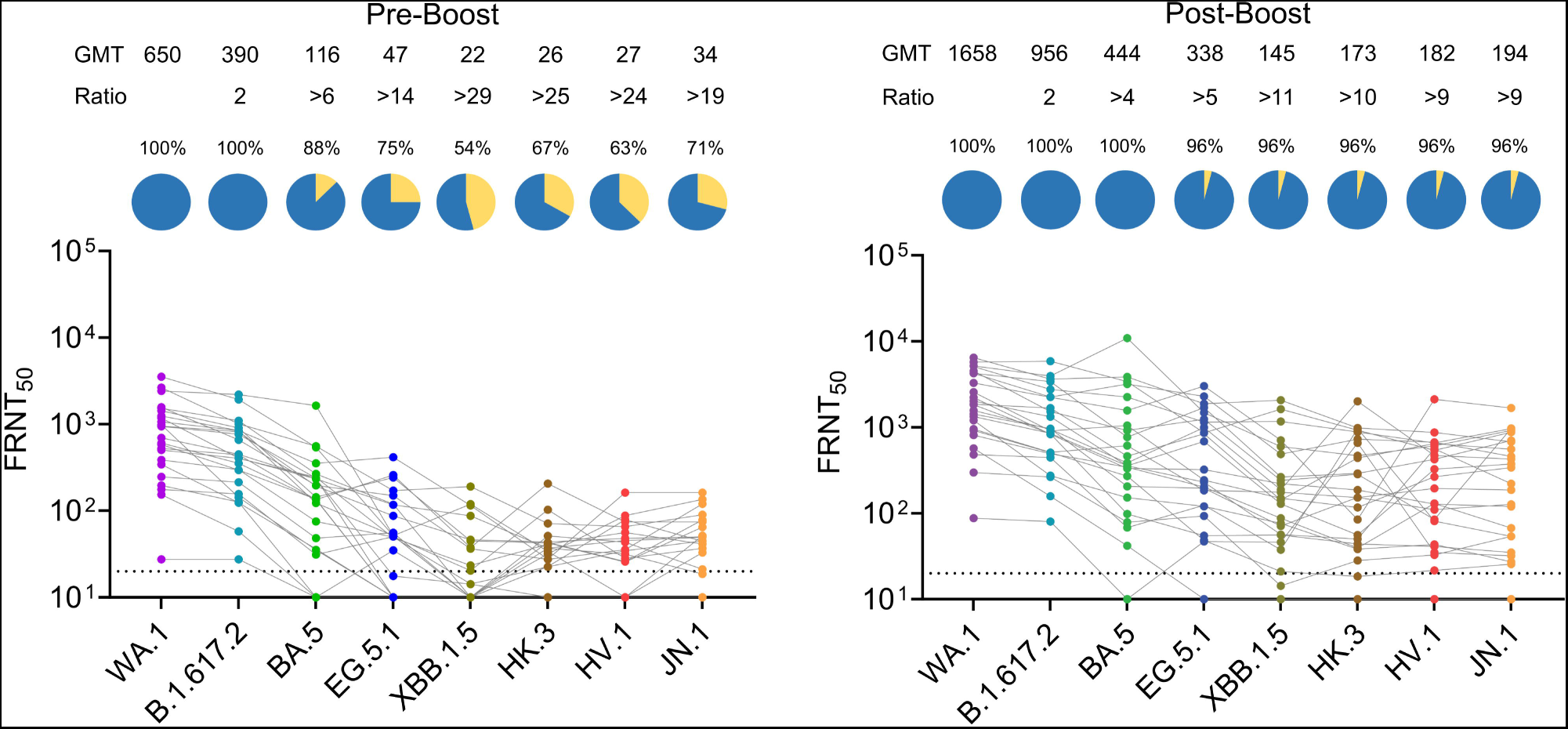
Live virus neutralization antibody responses against WA1/2020, B.1.617.2 and Omicron subvariants pre and post-XBB.1.5 vaccination. Shown is the neutralization activity against the WA1/2020 strain of severe acute respiratory syndrome coronavirus 2 (SARS-CoV-2), B.1.617.2 (Delta) variant and the Omicron subvariants BA.5, EG.5.1, XBB.1.5, HK.3, HV.1, and JN.1 in 24 participants (A) before and (B) after receiving XBB.1.5 booster immunization. The focus reduction neutralization test (FRNT_50_ [the reciprocal dilution of serum that neutralizes 50% of the input virus]) geometric mean titer (GMT) of neutralizing antibodies against WA1, B.1.617.2 variant and each Omicron subvariant are shown at the top of each panel, along with the ratio of the neutralization GMT against WA1 to that against B.1.617.2 and Omicron subvariants. The connecting lines between the variants represent matched serum samples. The horizontal dotted lines represent the limit of detection (LOD) of the assay (FRNT50 GMT 20). A pie chart above each graph shows the percentage of individuals who have titers above (responders) the LOD.

Next, we evaluated the neutralization response in the serum samples of all the individuals after XBB.1.5 monovalent booster. Importantly, the neutralizing antibody response against all the Omicron variants increased significantly after the XBB.1.5 booster in all the participants except one. After receiving the XBB.1.5 monovalent booster, the FRNT_50_ GMTs were 1658 against WA1, 956 against B.1.617.2, 444 against BA.5, 338 against EG.5.1, 145 against XBB.1.5, 173 against HK.3, 182 against HV.1, and 194 against JN.1. The neutralizing antibody titers against WA1 and B.1.617.2 in individuals after receiving the booster were 2.5-fold higher than neutralization titers before receiving the booster. After 13-37 days of the XBB.1.5 monovalent booster, all individuals except one (96%) had detectable neutralizing antibody titers against all the emerging Omicron variants, and the nAB titers were 4-7 higher than before the administration of vaccination (**Fig. 1b**).

### XBB.1.5 monovalent booster improves spike IgG binding to XBB.1.5 as compared to WA1

We next determined the total IgG and IgG subclass binding titers using spike-specific electro-chemiluminescence assays against ancestral SARS-CoV-2 and various Omicron variants. Nearly all samples before vaccination were positive for ancestral-spike total IgG binding (**Fig. 2a**). Prior to the XBB1.5 monovalent booster, IgG binding to the XBB.1.5 spike protein was lower by 6.1-fold (GMT: 32429; range: 5836-198765) as compared to WA1 spike protein (GMT: 198698; range: 30055–841492). In a similar manner, IgG binding to the BA.2.75 (4.8-fold), BA.1 (4.8-fold), BA.5 (3.4-fold), BQ.1.1 (5.6-fold) and XBB.1 (6.2-fold) spike proteins were lower as compared to WA1.

**Figure 2.**
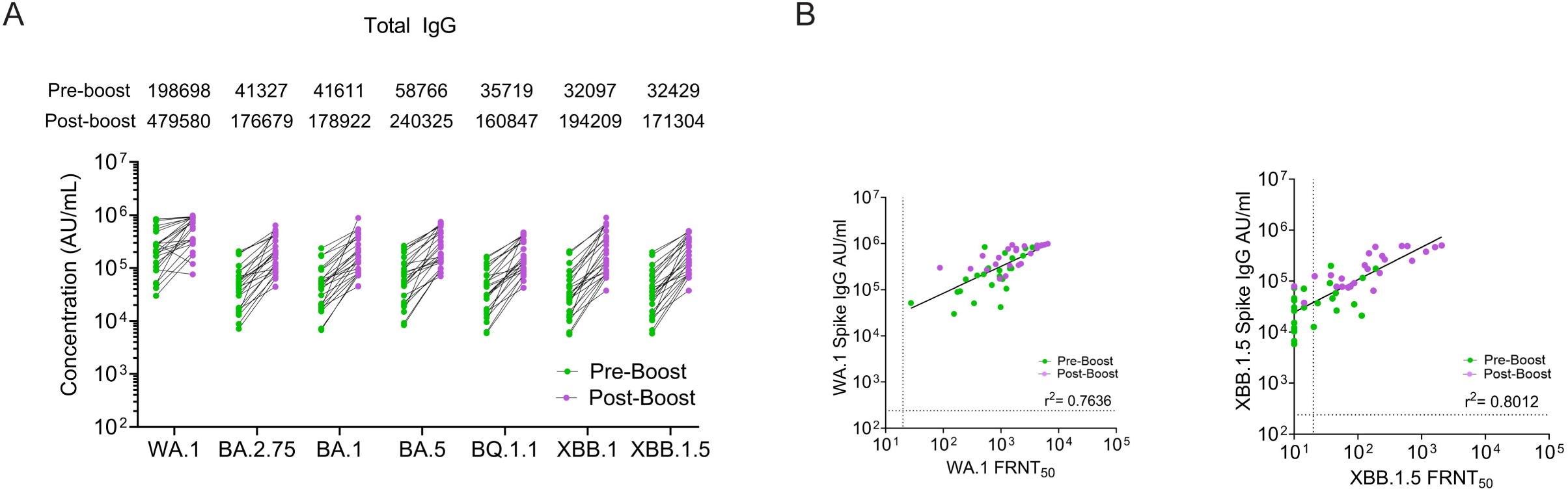
SARS-CoV-2 IgG spike binding following the XBB.1.5 moovalent booster dose. (**A**) Total IgG antibody responses against SARS-CoV-2 spike protein were measured by an electro-chemiluminescent multiplex immunoassay using the MSD platform and reported as arbitrary units per ml (AU/mL) as normalized by a standard curve for WA1 and Omicron subvariants BA.2.75. BA.1, BA.5, BQ.1.1, XBB.1, and XBB.1.5. Shown is the spike-binding total IgG antibody titers against the WA1 and the Omicron subvariants for the 24 participants before (green symbols) and after (pink symbols) receiving XBB.1.5 booster immunization. (**B**) Spearman correlation plots between the corresponding SARS-CoV-2 spike binding and FRNT_50_ for the WA1 and XBB.1.5 Omicron variant. The correlation plots between the corresponding spike and FRNT_50_ for the WA1 (panel B) and XBB.1.5 (panel C) variant are shown for the 24 individuals pre (green symbols) and post (pink symbols) XBB.1.5 booster vaccination. The normality of the antibody binding and neutralization titers was determined using a Shapiro-Wilk normality test. A nonparametric pairwise analysis for spike-specific IgG titers and neutralization titers was performed by a Wilcoxon matched-pairs signed rank test. A Spearman rank test was used for the correlation analysis of the spike-specific IgG AU/mL values against FRNT_50_ titers.

Following the XBB.1.5 monovalent booster, IgG binding to the cognate XBB.1.5 and related XBB.1 spike protein showed a greater increase (5.3-fold and 6.1-fold, respectively) as compared to binding to the WA1 spike protein (2.4-fold). In a similar manner, IgG binding to the BA.2.75 (4.3-fold) BA.1 (4.3-fold), BA.5 (4.1-fold), and BQ.1.1 (4.5-fold) spike protein were increased as compared to binding to the WA1 spike protein. For both pre- and post-boost, we observed a strong correlation between the corresponding spike-specific total IgG titers to WA1 neutralization titers (r^2^ = 0.6069; **Fig. 2b**) and the XBB.1.5 variant neutralization titers (r^2^ = 0.5436).

We next evaluated IgG subclass binding to the WA1, XBB.1.5 and related Omicron spike proteins. Prior to the XBB.1.5 monovalent booster, we observed IgG1 (GMT: 12.87; range: 0.52-114), IgG2 (GMT: 0.459; range: 0.05–6.27), IgG3 (GMT: 0.109; range: 0.04–5.85), and IgG4 (GMT: 3.346; range: 0.05–70.64) binding to the WA1 spike protein. In comparison, IgG subclass binding to the XBB.1.5 spike protein was substantially lower for IgG1 (GMT: 2.192; range: 0.07-24), IgG2 (GMT: 0.459; range: 0.02–0.81), IgG3 (GMT: 0.103; range: 0.01–0.18), and IgG4 (GMT: 0.057; range: <0.00006–24).

Following the XBB.1.5 monovalent booster, IgG1, IgG2, IgG3, and IgG4 binding to the WA1 spike protein increased by 1.8-fold, 2.4-fold, 3-fold, 3.2-fold, respectively (**Fig. 3**). On the other hand, IgG1, IgG2, and IgG3 binding to the XBB.1.5 spike increased by 3.7-fold, 4-fold, and 8.6-fold, respectively. In the case of IgG4 subclass antibodies, we observed no detectable IgG4 binding titer against the XBB.1.5 spike in 7 out of 24 participants (29%) before the booster immunization. In these 7 individuals, after the administration of XBB.1.5 monovalent booster, IgG4 binding titers (GMT: 0.22; range: 0.05-5.86) increased robustly against the XBB.1.5 spike protein. For the other 17 individuals, we observed an approximately 7-fold increase in binding to the XBB.1.5 spike after the XBB.1.5 booster dose. Most of the other Omicron subvariants, except for 2 individuals for XBB.1, showed pre-boost spike binding titers and were increased following the XBB.1.5 monovalent booster dose. Collectively, these data demonstrate that the XBB.1.5 monovalent booster results in a greater increase in spike binding to the cognate XBB.1.5 spike protein as compared to the WA1 spike protein. Further, these data demonstrate that all individuals post-boost show a more substantial increase of IgG4 against the XBB.1.5 spike protein.

**Figure 3.**
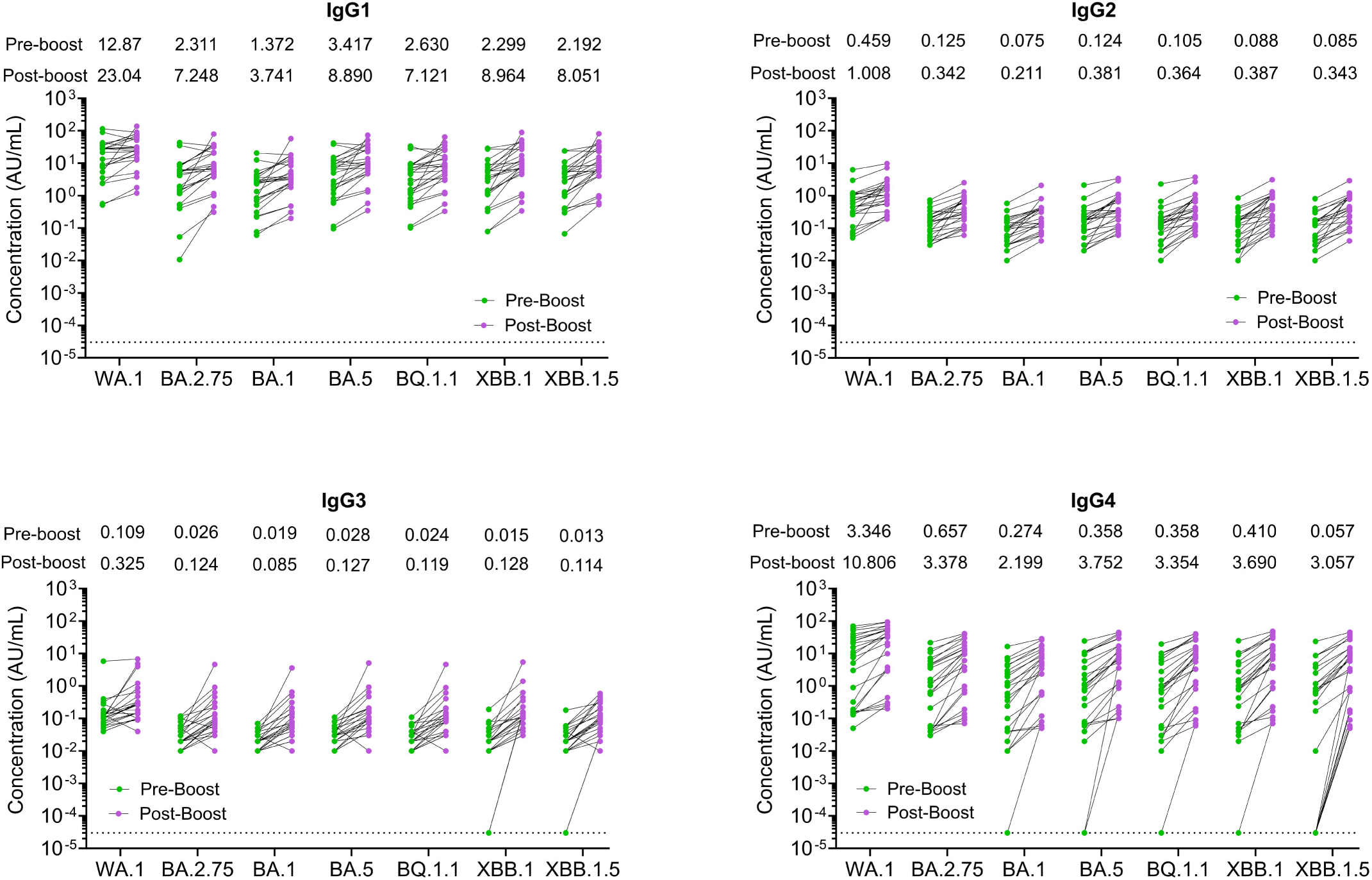
SARS-CoV-2 IgG subclass binding to SARS-CoV-2 variants following the XBB.1.5 moovalent booster dose. IgG antibody subclasses responses against SARS-CoV-2 spike proteins were measured by an electro-chemiluminescent multiplex immunoassay using the MSD platform and reported as arbitrary units per ml (AU/mL) as normalized by a standard curve using a monoclonal antibody. Shown are the spike-binding IgG subclasses antibody titers against the WA1 and the Omicron subvariants for the 24 participants before (green symbols) and after (pink symbols) receiving XBB.1.5 booster immunization. The dotted line in the spike binding assays represents the limit of detection.

### XBB.1.5 monovalent booster shows greater cross-reactive memory B cells that recognize epitopes outside of the RBD

We next determined how the XBB.1.5 monovalent booster impacts the frequency of spike-specific MBCs. We isolated PBMCs from the corresponding samples and performed flow cytometry on MBCs (CD3-CD14-CD19+CD20+ CD27+IgD-) using WA1 or XBB.1.5 trimeric spike (**Fig. 4a**) or RBD (**Fig. 4b**) probes directly conjugated to either AF488 (WA1) or BV421 (XBB.1.5). Prior to the XBB.1.5 booster, we observed 2.26% of MBCs were spike-specific, corresponding to 1.1% WA1+ only, 0.35% XBB.1.5+ only and 0.8% WA1+/XBB.1.5+ double - positive cells. In contrast, while the vast majority of the RBD-specific MBCs (2.03%) were WA1 RBD+ (1.91%), we observed very few XBB.1.5+ or WA1+/XBB.1.5+ MBCs (0.06% and 0.02%, respectively). Following the booster dose, we observed a substantial increase in cross-reactive-spike-specific MBCs as compared to cross-reactive RBD-specific MBCs. While we observed no increase in WA1+ spike MBCs, we observed an increase in XBB.1.5+ (0.46%) and WA1+/XBB.1.5+ (1.52%) spike-specific MBCs. In contrast, we observed a marginal increase in WA1+ (pre-boost: 1.91%; post-boost: 2.33%) and no increase in XBB.1.5+ RBD-specific MBCs (pre-boost: 0.06%; post-boost: 0.06%) and a small increase in WA1+/XBB.1.5+ RBD-specific MBCs (pre-boost: 0.02%; post-boost: 0.06%). These findings suggest that the XBB.1.5 monovalent booster primarily increases cross-reative antibodies located outside of the receptor binding region of the SARS-CoV-2 spike protein.

**Figure 4.**
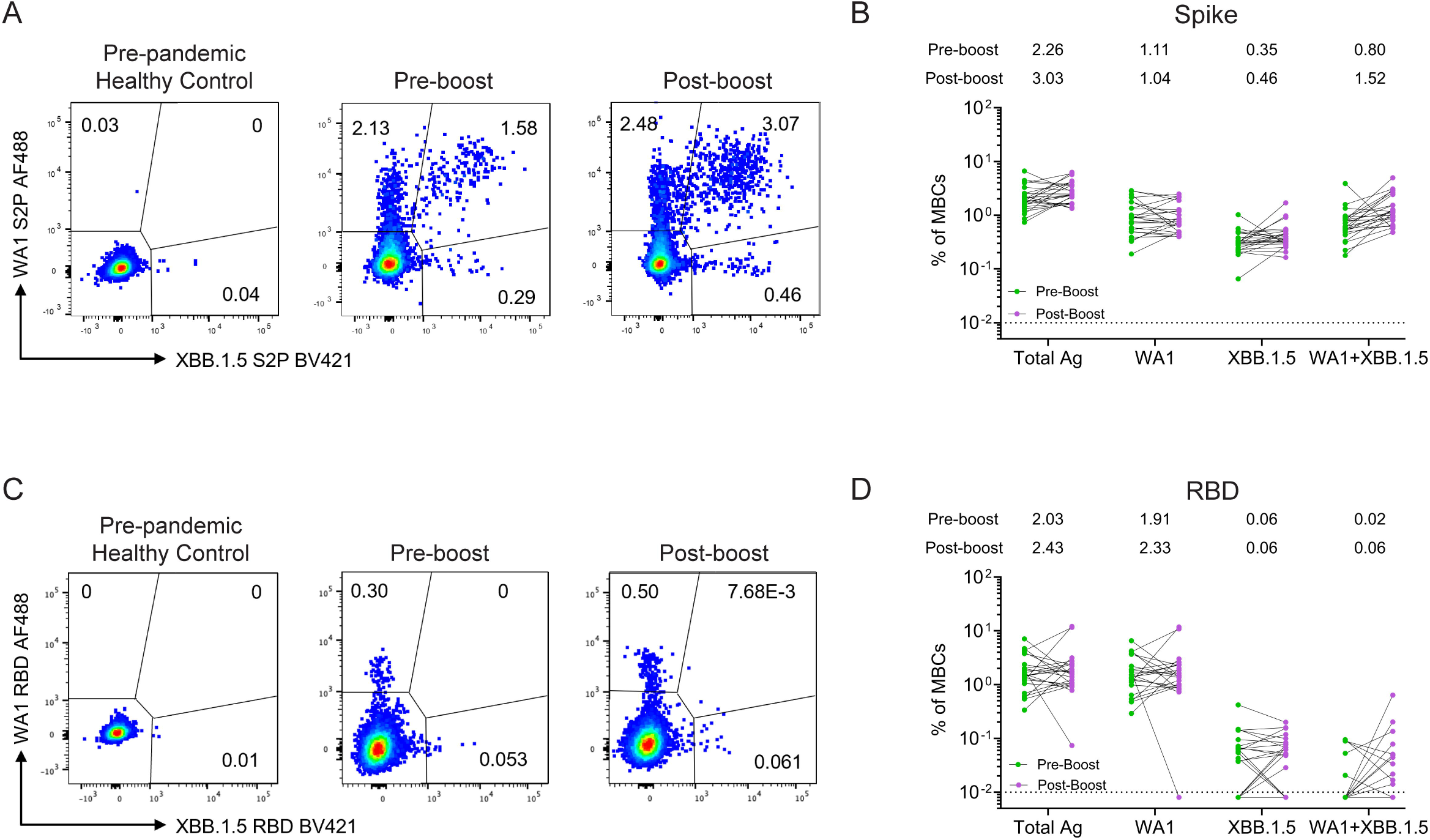
SARS-CoV-2 spike- and RBD-specific memory B cell responses following the XBB.1.5 moovalent booster dose. (A) Gating scheme of SARS-CoV-2 Spike S2P-specific MBCs, gated as live single CD19+IgD-CD20+CD27+ cells that are WA1 AF488+ and/or XBB.1.5 BV421+ cells, in a pre-pandemic healthy control, pre-XBB.1.5 booster and post-XBB.1.5 booster. **(B)** Spike S2P-specific MBCs are shown as the percentage of CD27+ MBCs in pre- and post-XBB.1.5 booster vaccinated individuals (n=24). The dotted line represents the percentage of Spike-specific MBCs in pre-pandemic healthy controls. **(C)** Gating scheme of SARS-CoV-2 RBD-specific MBCs, gated as live single CD19+IgD-CD20+CD27+ cells that are WA1 AF488+ and/or XBB.1.5 BV421+ cells, in pre-pandemic healthy controls, pre-XBB.1.5 booster and post-XBB.1.5 booster. **(D)** RBD-specific MBCs are shown as the percentage of CD27+ MBCs in pre- and post-XBB.1.5 booster vaccinated individuals (n=24). The dotted line represents the percentage of RBD-specific MBCs in pre-pandemic healthy control.

## Discussion

The findings in this study demonstrate that the antibodies elicited by the XBB.1.5 monovalent booster vaccine are effective at neutralizing the XBB.1.5 variant and other circulating Omicron subvariants, especially JN.1 and HV.1. Further, the serological data suggest that the XBB.1.5 monovalent booster overcomes immunological imprinting associated with the ancestral vaccines and SARS-CoV-2 infections. Consistent with this, we observed improved responses to XBB.1.5 spike compared to the WA1 spike protein as demonstrated by enhanced neutralizing activity, binding antibodies and memory B cells to XBB.1.5. These findings are comprable to prior studies evaluating neutralization breadth following the bivalent mRNA vaccine booster dose (WA1 and BA.5 spike)^8,14,15^.

We report that the XBB.1.5 monovalent mRNA booster results in a greater increase across all IgG subclasses, including IgG4. Early in the pandemic, high levels of anti-spike IgG4 antibodies were associated with poor clinical outcomes^16,17^. However, in cohorts of vaccinated healthcare workers, an increase in anti-spike IgG4 antibodies has been reported months after the second booster immunization with the ancestral monovalent mRNA vaccines^18^. The IgG4 class switching occurred after repeated mRNA vaccinations but not after viral vector-based immunization^19,20^. Preferential induction of IgG4 antibodies by vaccination has been reported to be associated with a high dose of vaccine or repeated immunization with the same antigen^21^ that resulted in the induction of specific T cell tolerance^22,23^. Interestingly, repeated immunization with a SARS-CoV-2 RBD-based vaccine induced tolerance in mice^24^. Further research is therefore needed to evaluate the mechanisms of the preferential IgG4 class switching induced by the XBB.1.5 monovalent mRNA vaccine, which is relevant for designing future seasonal SARS-CoV-2 booster immunizations.

It has been speculated that the emergence of isotype-switched memory B cells is consistent with a more robust and durable germinal center (GC) response induced by mRNA-based vaccines^25^. These findings contrast with the predominant induction of IgG1 and IgG3 after natural SARS-CoV-2 infections^26^. IgG4 has limited functional activity due to structural features that impair its binding to C1q and Fc receptors that mediate opsonization and antibody-depending cellular cytotoxicity^27^. These features likely make IgG4 a blocking or suppressing antibody^28^. Further investigations are therefore required to define the role of anti-spike IgG4 induced by XBB.1.5 monovalent booster vaccine in protection or its impact on the response to future booster immunizations

We have previously shown that Omicron infection induces a more balanced neutralizing antibody response to Omicron variants, supporting the the need to include an Omicron spike in vaccine formulations^29^. We and others also found that the bivalent WA1/BA.5 mRNA vaccines improved neutralizing activity and breadth against BA.5 and related Omicron variants^8,11,30–32^. However, the inclusion of the ancestral spike in vaccine formulations has raised concerns about immunological imprinting^33,34^. Thus, the FDA recently made the decision to reformulate mRNA vaccines with a single monovalent XBB.1.5 spike. In our studies, we probed MBCs before and after XBB.1.5 monovalent booster and found that a majority of cross-reactive spike-specific MBCs with marginal cross-reactivity within the RBD. This suggests that the XBB.1.5 monovalent booster is inducing non-RBD cross-reactive antibodies and hence backboosting antibodies against the ancestral spike. Further studies are needed to determine the non-RBD epitopes targeted following XBB.1.5 monovalent booster administration.

The limitations of this study include the small cohort size and short follow-up after booster dose, the lack of T cell analysis, and the unknown effect of previous exposure to SARS-CoV-2 infection(s). Future longitudinal studies with more individuals would be needed to address the prevalence of IgG4 class switching induced by repeated boosting with the mRNA vaccine. Further, immunological imprinting continues to be a concern as the XBB.1.5 monovalent booster is unable to induce a robust XBB.1.5 RBD-specific MBC response.

---

**Supplementary Figure 1.**
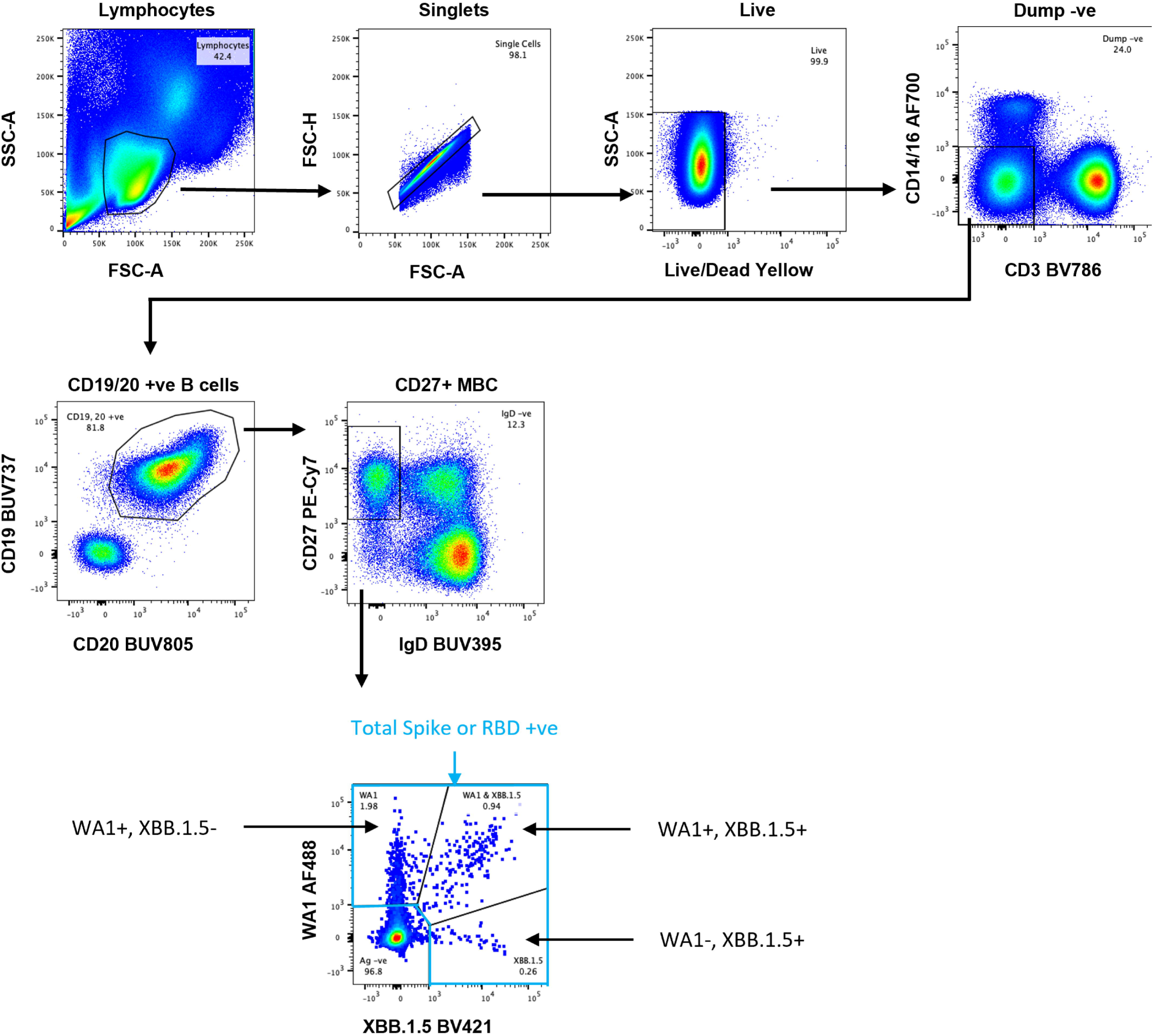
Gating strategy to identify SARS-CoV-2 Spike S2P or RBD-specific total, WA1 or XBB.1.5 or WA1 and XBB.1.5 dual positive cells.

## Author Contributions

S.J., A.M., and M.S.S. contributed to the analysis, interpretation of data and writing the manuscript. L.L. performed in vitro neutralization experiments. S.J., A.A.M. and R.W. processed the samples for PBMCs isolation. S.J. and M.L.E. performed binding assays. S.J. and S.K. contributed to the B cell experiments, acquisition, analysis and writing the manuscript. S.L. performed IgG subclass calculations and approval of the final manuscript. S.G., D.S., M.K.S., K.B., I.P., R.T., B.K., S.M.D., M.B.L., S.T.W., C.C., C.R., R.W., I.T.T., D.W., P.D.K., N.R., B.A.P., and D.C.D. provided clinical support for the study and contributed to sample collection. J.W., A.M., and M.S.S. contributed to the conception and design of the work and the writing and approval of the final manuscript.

## Methods

### Serum samples

The peripheral blood samples from donors were collected at Emory University (Emory National Primate Research Center, Emory Hope Clinic, and Emory Children’s Center). Adults ≥18 years old who met eligibility criteria under these protocols and provided informed consent were enrolled. All patient samples were de-identified prior to inclusion in the study. Collection and processing were performed under approval from the University Institutional Review Board (#00002061,#00058271 and #00022371). Of all the donors, samples were collected before (0-32 days) the XBB.1.5 monovalent booster and 13-41 days after XBB.1.5 booster. Blood samples were processed to obtain plasma and peripheral mononuclear cells.

### Cells and Viruses

VeroE6-TMPRSS2 cells were generated and cultured as previously described^35^. nCoV/USA_WA1/2020 (WA1), closely resembling the original Wuhan strain, was propagated from an infectious SARS-CoV-2 clone as previously described^36^. B.1617.2 (EPI_ISL_2457061) was provided by Dr. Richard Webby (St Jude Children’s Research Hospital). Omicron subvariants were isolated from residual nasal swabs. BA.5 isolate (EPI_ISL_13512579) was provided by Dr. Richard Webby and XBB.1.5 (EPI_ISL_16026423) was provided by Dr. Andrew Pekosz. EG.5.1 (EPI_ISL_17977757), HV.1 (EPI_ISL_18403105), HK.3 (EPI_ISL_18403093), and JN.1 (EPI_ISL_18403077) were provided by Dr. Benjamin Pinsky (Stanford University) (**Supplementary Table S2**). All variants were plaque purified and propagated once in VeroE6-TMPRSS2 cells to generate working stocks. Viruses were deep sequenced and confirmed as previously described^37^.

### Focus Reduction Neutralization Test (FRNT)

FRNT assays were performed as previously described^35,37,38^. Briefly, serum samples in duplicate were 3-fold diluted in 8 serial dilutions using DMEM with an initial dilution of 1:10. Serially diluted samples were mixed with an equal volume of SARS-CoV-2 (100-200 foci per well). The virus-serum mixtures were incubated at 37°C for 1 hour in a round bottom 96-well culture plate. After 1 hour incubation, the virus-serum mixture was added to VeroE6-TMPRSS2 cells and incubated at 37°C for an additional hour. Post-incubation, the mixture was removed from cells and 100μl of prewarmed 0.85% methylcellulose overlay was added to each well. Plates were incubated at 37°C for 18 to 40 hours (depending on variants). After the appropriate incubation time, the methylcellulose overlay was removed, and cells were washed with PBS and fixed with 2% paraformaldehyde for 30 minutes. Following fixation, cells were washed twice with PBS, and permeabilized using permeabilization buffer for at least 20 minutes. After permeabilization, cells were incubated with an anti-SARS-CoV-2 spike primary antibody directly conjugated to Alexa Fluor-647 (CR3022-AF647) overnight at 4°C. Cells were then washed twice with 1X PBS and imaged on an ELISPOT reader (CTL Analyzer).

### Spike-binding assay

SARS-CoV-2 spike-specific IgG antibodies and IgG subclasses were detected using electro chemiluminescent-based multiplex immunoassay on the Meso 47 Scale Discovery (MSD) platform using V-PLEX SARS-CoV-2 Key Variant Spike Panel 1 Kit (catalog 48 number K15651 (IgG)), Panel 34 (catalog number K15690 (IgG)), and in-house made IgG subclass antibodies. Briefly, serum samples were diluted 1:5000 prior to the assay. Kit plates were blocked for 30 minutes using the manufacturer-provided blocking buffer A. The blocking buffer was removed, and the plates were washed 3 times with wash buffer. Diluted serum samples, mesoscale standard controls, and SARS-CoV-2 spike-specific IgG subclass monoclonal antibody controls were added to the plates and incubated for 2 hours at room temperature on a plate shaker adjusted to 700 rpm. After a 2-hour incubation, plates were washed 3 times, loaded with a solution containing either MSD Sulfo-Tag anti-human IgG or mouse anti-human IgG subclass antibodies (mAbtech), and incubated for 1 hour at room temperature on a plate shaker at 700 rpm. For IgG subclass measurements, plates were washed 3 times and 25ul/well of 5ug/ml MSD Sulfo-tag anti-mouse antibody was added and incubated for an additional hour. Plates were subsequently washed 3 times and 150ul/well of detection read buffer was added immediately prior to the plate read. Raw data were analyzed using Discovery Workbench and GraphPad Prism software v10.1.2. Plasma antibody concentrations were calculated relative to antibody standard controls and expressed in arbitrary units (AU) per mL.

### S2P and RBD antigens

Biotinylated probes of the stabilized SARS CoV-2 spike trimer S2P and the RBD subdomain were produced as previously described^39,40^. Briefly, the DNA sequence encoding either the S2P or RBD region from ancestral WA1 strain or XBB.1.5 variant and flanked by an HRV3C cleavable single-chain Fc tag and an AVI tag was cloned in a mammalian expression vector and transfected in FreeStyle 293 cells. Six days post transfection, the tagged protein in culture supernatant was captured by protein A resin and collected in the flow-through after concurrent HRV3C cleavage and BirA biotinylation overnight. Finally, the biotinylated probe was purified on a Superdex 200 16/600 gel filtration column equilibrated with PBS.

### Multicolor immunophenotyping of B cells by flow cytometry

The peripheral blood mononuclear cells (PBMCs) were isolated from the collected blood samples of donors and cryopreserved in liquid nitrogen until use. For B-cell probe binding, the cryopreserved PBMCs were thawed at 37°C and resuspended in pre-warmed RPMI media supplemented with 10% Fetal Bovine Serum (FBS). The PBMCs were washed twice with FACS buffer (1% BSA in 1x PBS) and incubated with antigen probe mix containing streptavidin-AlexaFluor488 labeled WA1 S-2P or RBD and streptavidin-BV421 labeled XBB.1.5 S-2P or RBD for 60 minutes on ice and washed twice with FACS buffer. Following incubation, cells were stained with a panel of cell surface antigen specific antibodies (**Supplementary Table S3**), CD3 BV786, CD14/16 Alexa Fluor700, CD19 BUV737, CD20 BUV805, CD27 PE-Cy7, IgD BUV395, IgG1 BV650, IgG2 PE, IgG3 BV605, and IgG4 AlexaFluor594 for half an hour on ice. Cells were washed twice with 1x PBS and incubated with live/dead stain for 10 minutes on ice. After live/dead staining, cells were washed twice and resuspended in FACS buffer for acquisition using FACSymphony A5. The data were analyzed using FlowJo version 10.9.0 (BD Bioscience).

### Quantification and Statistical Analysis

Antibody neutralization was quantified by counting the number of foci for each sample using the Viridot program^41^. The neutralization titers were calculated as follows: 1 – (ratio of the mean number of foci in the presence of sera and foci at the highest dilution of the respective sera sample). Each sample was tested in duplicate. The FRNT-50 titers were interpolated using a 4-parameter nonlinear regression in GraphPad Prism 10.1.2. Samples that do not neutralize at the limit of detection at 50% are plotted at 20 and used for geometric mean and fold-change calculations. The normality of the antibody binding and neutralization titers were determined using the Shapiro-Wilk normality test. Nonparametric pairwise analysis for spike-specific IgG titers and neutralization titers were performed by Wilcoxon matched-pairs signed rank test. Spearman rank test was used for the correlation analysis. Correlations analyses were done by log transforming spike binding titers or neutralization titers, followed by linear regression analysis. The r^2^ and *p* values are reported in each figure.

