## Supplementary Text for "XBB.1.5 monovalent booster improves antibody binding and neutralization against emerging SARS-CoV-2 Omicron variants"

**Supplementary Table S1: Demographic information for individuals in the study group**

| **General information** | | | **Prior vaccination history** | | | | **COVID history** | | **Days from XBB.1.5 vaccination** |
| --- | --- | --- | --- | --- | --- | --- | --- | --- | --- |
| **Sample** | **Age** | **Sex** | **Primary Vaccination 1** | **Primary Vaccination 2** | **Monovalent Booster** | **Bivalent**  **Booster** | **SARS-COV-2 infection** | **If yes, Date** |  |
| 1 | 70 | F | Moderna | Moderna | Moderna | 28-Sep-22 | No |  | 13 |
| 2 | 41 | F | Moderna | Moderna | Moderna |  | No |  | 29 |
| 3 | 35 | F | Moderna 01/02/21 | Moderna 1/30/21 | Moderna 10/23/21 | Moderna 17-Sep-22 | no |  | 24 |
| 4 | 28 | M | Pfizer 22-Jan-21 | Pfizer 12-Feb-21 | Pfizer 6-Jan-22 |  | Yes | 12-Jun-22 | 22 |
| 5 | 27 | M | Moderna 15-Jan-21 | Moderna 15-Feb-21 | Pfizer 15-Nov-22 |  | Yes | 21-Jun-22 | 28 |
| 6 | 27 | M | Pfizer 19-Jan-21 | Pfizer 10-Feb-21 | Pfizer 6-Dec-21 |  | N/A |  | 29 |
| 7 | 34 | F | Pfizer 4-Jan-21 | Pfizer 22-Jan-21 | Pfizer 30-Aug-21 | Pfizer 7-Sep-22 | No |  | 27 |
| 8 | 42 | F | Moderna 8-Jan-21 | Moderna 6-Feb-21 | Moderna 27-Dec-21 |  | N/A |  | 24 |
| 9 | 25 | M | Pfizer 5-May-21 | Pfizer 27-May-21 |  |  | Yes | 16-Jan-23 | 18 |
| 10 | 31 | M | Pfizer 1-Apr-21 | Pfizer 22-Apr-21 |  |  | Yes | 1/1/2022, 6-Jun22 | 18 |
| 11 | 25 | F | Pfizer 30-Mar-21 | Pfizer 20-Apr-21 | Pfizer 10-Nov-21 |  | Yes | 15-Aug-22, 15-mar-23 | 22 |
| 12 | 21 | F | NVX-COV2373 15-Jan-21 | NVX-COV2373 15-Feb-21 | Moderna 15-Nov-21 |  | Yes | 15-Mar-20 | 26 |
| 13 | 57 | M | 23-Mar-21 | 20-Apr-21 | 30-Nov-21 | 28-Nov-22 | Yes | 13-Mar-20, 5/15/2022 | 31 |
| 14 | 33 | F | 4-Feb-21 | 2-Mar-21 | 5-Dec-21 | 3-Dec-22 | Yes | 6/21/2022, 8/11/2022 | 33 |
| 15 | 32 | M | 31-Mar-21 | 30-Apr-21 | 12-Jan-22 | 11-Nov-22 | Yes | 31-Dec-22 | 26 |
| 16 | 39 | M | 20-Mar-21 | 10-Apr-21 | 16-Oct-21 | 12-Nov-22 | Yes | 3-Nov-20 | 37 |
| 17 | 31 | M | 1-Aug-21 | 3-Feb-21 | 20-Dec-21 | 29-Oct-22 | No |  | 28 |
| 18 | 26 | M | 30-Dec-20 | 18-Jan-21 | 29-Aug-21 | 7-Sep-22 | No |  | 34 |
| 19 | 45 | M | 20-Mar-21 | 10-Apr-21 | 16-Oct-21 | 12-Nov-22 | Yes | 11/1/2020, 5/1/22 | 35 |
| 20 | 24 | M | 10-May-21 | 1-Jun-21 | 22-Dec-21 | 17-Oct-22 | Yes | 1-Jan-22 | 41 |
| 21 | 61 | M | 7-Jan-21 | 4-Feb-21 | 28-Oct-21 | 22-Oct-22 | Yes | 11/3/2020, 6/1/22 | 33 |
| 22 | 68 | M | 13-Jan-21 | 10-Feb-21 | 4-Nov-21 | 17-Oct-22 | Yes | 22-Nov-22 | 28 |
| 23 | 21 | F | 31-Mar-21 | 28-Apr-21 | 7-Dec-21 | 20-Nov-22 | Yes | 27-Jan-22 | 31 |
| 24 | 56 | F | 20-Aug-20 | 21-Sep-20 | 19-Nov-21 | 17-Oct-22 | Yes | 15-Jul-22 | 28 |

**Supplementary Table S2: Amino acid substitutions in spike protein of variants used in the study**

| **Variant** | **Amino acid substitutions** | **Virus Name** | **GISAID** |
| --- | --- | --- | --- |
| B.1617.2 | T19R, G142D, N282S, **L452R, T478K,** D614G, P681R, D950N | hCoV-19/USA/CA-Stanford-24_S29/2021 | EPI_ISL_2457061 |
| BA.5 | T19I, L24del, P25del, P26del, A27S, H69del, V70del, T76I, G142D, V213G, **G339D, S371F, S373P, S375F, T376A, D405N, R408S, K417N, N440K, L452R, S477N, T478K, E484A, F486V, Q498R, N501Y, Y505H,** D614G, H655Y, N679K, P681H, N764K, D796Y, Q954H, N969K | hcov- 19/USA/MD/HP30386/202 2 | EPI_ISL_1351257 9 |
| XBB.1.5 | T19I, L24del, P25del, P26del, A27S, V83A, G142D, Y144del, H146Q, Q183E, V213E, G252V, **G339H, R346T, L368I, S371F, S373P, S375F, T376A, D405N, R408S, K417N, N440K, V445P, G446S, N460K, S477N, T478K, E484A, F486P, F490S, Q498R, N501Y, Y505H,** D614G, H655Y, N679K, P681H, N764K, D796Y, N969K, Q954H | hCoV-19/USA/MD-HP40900-PIDYSWHNUB/2022 | EPI_ISL_16026423 |
| EG.5.1 | T19I, L24del, P25del, P26del, A27S, Q52H, V83A, G142D, Y144del, H146Q, Q183E, V213E, G252V, **G339H, R346T, L368I, S371F, S373P, S375F, T376A, D405N, R408S, K417N, N440K, V445P, G446S, F456L, N460K, S477N, T478K, E484A, F486P, F490S, Q498R, N501Y, Y505H,** D614G, H655Y, N679K, P681H, N764K, D796Y, N969K, Q954H | hCoV-19/USA/CA-Stanford-147_S01/2023 | EPI_ISL_17977757 |
| HK.3 | T19I, L24del, P25del, P26del, A27S, Q52H, V83A, G142D, Y144del, H146Q, Q183E, V213E, G252V, **G339H, R346T, L368I, S371F, S373P, S375F, T376A, D405N, R408S, K417N, N440K, V445P, G446S, L455F, F456L, N460K, S477N, T478K, G482R, E484A, F486P, F490S, Q498R, N501Y, Y505H,** D614G, H655Y, N679K, P681H, N764K, D796Y, N969K, Q954H | hCoV-19/USA/CA-Stanford-165_S29/2023 | EPI_ISL_18403093 |
| HV.1 | T19I, L24del, P25del, P26del, A27S, Q52H, V83A, G142D, Y144del, H146Q, F157L, Q183E, V213E, G252V, **G339H, R346T, L368I, S371F, S373P, S375F, T376A, D405N, R408S, K417N, N440K, V445P, G446S, L452R, F456L, N460K, S477N, T478K, E484A, F486P, F490S, Q498R, N501Y, Y505H, Y505H,** D614G, H655Y, N679K, P681H, N764K, D796Y, N969K, Q954H | hCoV-19/USA/CA-Stanford-165_S45/2023 | EPI_ISL_18403105 |
| JN.1 | T19I, R21T, L24del, P25del, P26del, A27S,S50L, H69del, V70del, V127F, G142D, Y144del, F157S, R158G, N211del, L212I, V213G, L216F, H245N,A264D, **I332V, G339H, K356T, S371F, S373P, S375F, T376A, R403K, D405N, R408S, K417N N440K, V445H, G446S, N450D, L452W, L455S, N460K, S477N, T478K, N481K, V483del, E484K, F486P, Q498R, N501Y, Y505H,** E554K, A570V, D614G, H655Y, N679K, P681R, N764K, D796Y, S939F, Q954H, N969K, P1143L, ins16MPLF | hCoV-19/USA/CA-Stanford-165_S10/2023 | EPI_ISL_18403077 |

**Bold letter indicates mutations in RBD region*

| **Target** | **Clone** | **Catalog No.** | **Fluorophore** | **Manufacturer** |
| --- | --- | --- | --- | --- |
| CD3 | SK7 | 563799 | BV786 | BD |
| CD14 | 61D3 | 56-0149-42 | AF700 | eBioscience |
| CD16 | eBioCB16 | 56-0168-42 | AF700 | BD |
| CD19 | SJ25C1 | 612756 | BUV737 | BD |
| CD20 | 2H7 | 612905 | BUV805 | BD |
| CD27 | O323 | 302838 | PE-Cy7 | Biolegend |
| IgD | IA6-2 | 563813 | BUV395 | BD |
| IgM | G20-127 | 551062 | BV605 | BD |
| IgA | IS11-8E10 | 130-113-472 | APC | BD |
| SA-AF488 |  | S32354 | AF488 | Invitrogen |
| SA-BV421 |  | 563259 | BV421 | BD |

**Supplementary Table S3: Antibody Panel**
